# Meta-Analysis of Urinary Metabolite GWAS Studies Identifies Three Novel Genome-Wide Significant Loci

**DOI:** 10.1101/2024.06.25.600593

**Authors:** Jihan K. Zaki, Jakub Tomasik, Jade McCune, Oren Scherman, Sabine Bahn

## Abstract

Genome-wide association studies (GWAS) have substantially enhanced the understanding of genetic influences on phenotypic outcomes; however, realizing their full potential requires an aggregate analysis of numerous studies. Here we represent the first comprehensive meta-analysis of urinary metabolite GWAS studies, aiming to consolidate existing data on metabolite-SNP associations, evaluate consistency across studies, and unravel novel genetic links. Following an extensive literature review and data collection through the EMBL-EBI GWAS Catalog, PubMed, and metabolomix.com, we employed a sample size-based meta-analytic approach to evaluate the significance of previously reported GWAS associations. Our results showed that 1226 SNPs are significantly associated with urinary metabolite levels, including 48 lead SNPs correlated with 14 analytes: alanine, 3-aminoisobutyrate, betaine, creatine, creatinine, formate, glycine, glycolate, histidine, 2-hydroxybutyrate, lysine, threonine, trimethylamine, and tyrosine. Notably, the results revealed three previously unknown associations: rs4594899 with tyrosine (P = 6.6 × 10^-9^, N = 5447), rs1755609 with glycine (P = 3.3 × 10^-10^, N = 5447), and rs79053399 with 3-aminoisobutyrate (P = 6.9 × 10^-10^, N = 4656). These findings underscore the potential of urinary metabolite GWAS meta-analyses in revealing novel genetic factors that may aid in the understanding of disease processes, and highlight the necessity for larger and more comprehensive future studies.

**Author summary:** In this extensive study, we’ve meta-analysed data from various genome-wide association studies to better understand the genetic determinants of urinary metabolite levels. These metabolites are small molecules found in urine that reflect our body’s biochemical activities and can indicate states of health or disease. By combining results from previous research, we’ve identified 48 significant independent associations between single nucleotide polymorphisms (SNPs) and the levels of 14 metabolites in urine. Among these, we highlight three novel SNP-metabolite associations that offer new insights into the genetic architecture underlying metabolite regulation.

Our findings contribute to the growing body of evidence that demonstrates the value of large-scale genetic meta-analyses. The significant SNP-metabolite links uncovered may serve as biomarkers for complex biological processes and disease mechanisms. This work takes us a step closer to a more nuanced understanding of the genetic factors that influence metabolic pathways and holds promise for improving diagnostic and therapeutic strategies through precision medicine. However, the complexity of genetic contributions to metabolite variations calls for continued research, underscoring the need for larger and more comprehensive studies in the future.

## Introduction

The identification of peripheral biomarkers holds significant promise for uncovering the mechanisms of complex diseases, improving diagnostic accuracy, and enabling prognosis and personalized treatment. One way to identify biomarkers is through genome-wide association studies (GWASs), which have transformed our knowledge of how genetic differences, typically in the form of single nucleotide polymorphisms (SNP), affect phenotypic outcomes [1]. While most initial GWAS studies focused on identifying genetic variants associated with specific binary traits or disorders [1], there has been a subsequent increase in focus towards more quantitative outcomes, such as biological analyte concentrations. The first metabolite GWAS study, conducted in 2008 [2], demonstrated the potential to establish robust causal links between genetic risk factors and target metabolites, and emphasized the importance of identifying relevant molecular intermediaries for a more complete understanding of the mechanistic pathways of disease. This argument has become stronger over the past decade, as the assessment of causal inference of metabolite levels on various diseases has become more accessible due to advancements in Mendelian randomization (MR) methodology and analyte measurement methods.

While numerous metabolite GWAS reports have been published since the first study, these have focused primarily on serum and plasma metabolites as opposed to other sample types, such as urine. This trend is exemplified by the fact that a PubMed search for serum and plasma GWAS studies currently returns nearly twenty times more results than for urine-based studies. Nonetheless, since the first urinary metabolite GWAS study was published in 2011 [3], research in this area has been steadily expanding, even though the overall sample sizes have remained relatively similar. In the face of limited large-scale analyses, a potential way to bolster the power of urinary metabolite GWAS studies is through the implementation of meta-analytic methods [4]. The combination of results from multiple studies could help uncover novel associations and strengthen confidence in previously reported findings.

In the present study, we aimed to conduct a comprehensive meta-analysis of existing urinary metabolite GWAS studies to combine all known metabolite-SNP associations into a single resource, assess the consistency of associations between studies, and identify novel genetic associations to urinary metabolite levels. A comprehensive database search was conducted to identify published and proprietary urinary metabolite GWAS data, followed by a sample size-based meta-analysis to identify significant genetic associations.

## Methods

### Metabolite data collection

The present meta-analysis included GWAS data obtained from three different sources, including the EMBL-EBI GWAS catalog under the trait urinary metabolite measurement (EFO-0005116); urinary metabolite GWAS studies compiled by Professor Karsten Suhre on metabolomix.com; and the PubMed library using the search term: ((GWAS) OR (genome-wide association) OR (Mendelian randomization)) AND (urine) AND ((metabolite) OR (analyte) OR (measurement) OR (biomarker)). Studies were considered eligible provided they included healthy participants, or did not study known specific metabolite-disorder associations, and had no overlapping participants with other studies. In cases where the GWAS summary statistic data were not publicly available, a data request was sent to the corresponding author(s), followed by a reminder. Only those metabolites that appeared in two or more studies, with two or more overlapping SNPs, were included in the meta-analysis. Metabolites whose identities were uncertain, due to the limitations of untargeted identification methods, were excluded to increase the accuracy of compound identification.

### Data processing

Information was extracted from the obtained GWAS summary statistics data regarding the measured analyte name, chromosome, position, the reference SNP cluster identifier (rsID) tag, effect allele, minor allele, beta coefficient, standard error, p-value, sample size, and effect allele frequency. Each evaluated analyte-SNP combination was harmonized to ensure allele consistency, i.e. SNPs with opposite alleles were flipped by changing the direction of the effect sizes as well as allele frequencies, and SNPs with non-matching effect alleles following flipping were removed.

### Sample size-based meta-analysis

The present analysis was conducted using R version 4.2.0. The meta-analysis utilized a sample size-weighted approach [5]. This method offers substantial flexibility, allowing for the integration of results even in the absence of effect size estimates or when the beta coefficients and standard errors from individual studies are not measured on the same scale. It was selected as the primary analysis method because most of the metabolomic GWAS data in the literature have incomparable effect sizes and standard errors, attributable to inconsistent measurement methods and inconsistent data transformation practices. The method applies Equations 1 and 2 to identify a combined Z-score from combined SNP summary statistics, which is then used to calculate the new, composite P-values. In brief, the method applies a weighted sum on calculated Z-scores from each individual study to identify a combined Z-score, where higher Z-scores correspond to lower P-values, and vice versa.

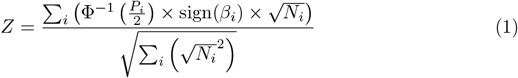

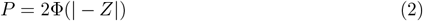

Where:

- *β*_*i*_: Effect size estimate of study
- Φ^*−*1^: Quantile function
- Φ: Distribution function
- *N*_*i*_: Sample size of study
- *P* : P-value
- *Z*: Z-score

### Post-processing and post-hoc analyses

A Bonferroni-corrected significance threshold of P < 7.1 × 10^-9^ was applied to adjust for multiple comparisons. To identify the final set of lead SNPs among the significant SNPs, all closely positioned (within 1000 kb) and correlated (r^2^ > 0.1) significant associations were combined using clumping, with the European population from the 1000 Genomes Project used as a reference panel [6]. A Manhattan plot of all evaluated associations was generated using the CMplot package in R. Potential confounding effects were assessed by reviewing the respective literature and searching the PhenoScanner database [7, 8]. Additionally, a pathway analysis was conducted for metabolites with significant associations clustered in nearby chromosomal regions using the MetaboAnalyst platform [9]. The hypergeometric test was applied as the enrichment evaluation method, and the analysis was based on the Kyoto Encyclopedia of Genes and Genomes Homo sapiens reference pathway library.

## Results

The current study consisted of a comprehensive literature review of urinary metabolite GWAS studies carried out to date, followed by extensive data collection and subsequent meta-analysis. Of the identified 191 unique studies of interest, 26 satisfied the inclusion criteria concerning the required urinary metabolite GWAS data and its quality. Following data enquiries where GWAS summary statistics had not been made publicly available, five studies were included in the final, sample size-based meta-analysis. A summary of the data collection process alongside the list of final included studies is shown in Figure 1.

**Fig 1.**
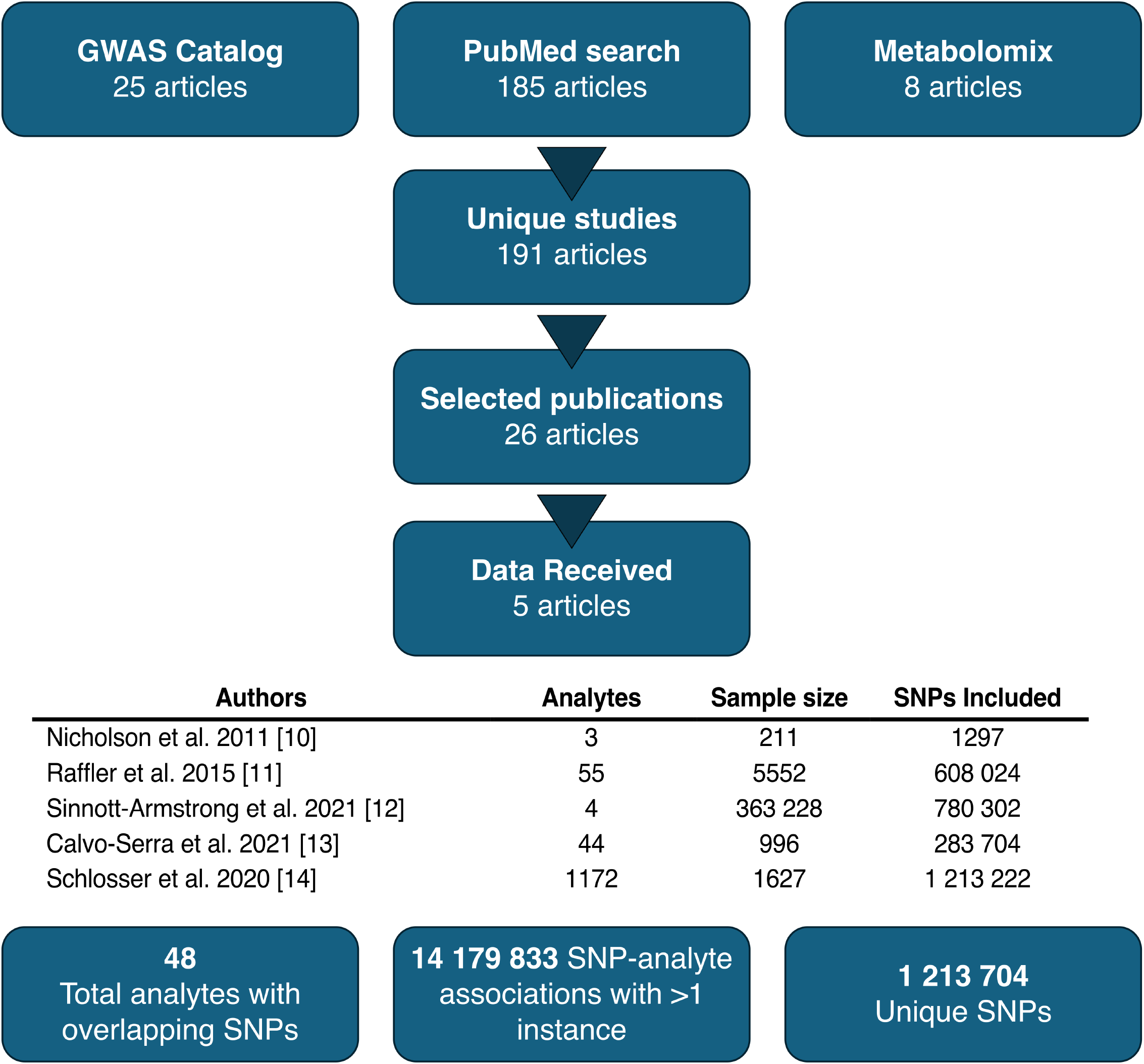
A summary of the data collection process for the urinary metabolite GWAS meta-analysis.

### Description of the included studies

A total of five studies were included in the meta-analysis, as summarized in Supplementary Table S1. The first study, by Nicholson et al. [10], aimed to identify genetic associations for three urinary analytes in two cohorts, MolTWIN and MolOBB, comprising 211 participants. Urinary metabolites were measured using ^1^H NMR and genotyping was performed using the Illumina 317K BeadChip SNP array. In the second study, Raffler et al. [11] measured 55 urinary markers using targeted NMR measurements in 3861 participants from the Study of Health in Pomerania (SHIP) and 1691 participants from the Kooperative Gesundheitsforschung in der Region Augsburg (KORA) study. Both analyses used Affymetrix Human SNP Array 6.0 gene chips for genotyping. Sinnott-Armstrong et al. [12] conducted a GWAS study of 4 urine analytes measured using clinical tests in 363 228 UK Biobank participants. Genotyping was conducted using the UK Biobank Axiom Array. Calvo-Serra et al. [13] performed a GWAS study aiming to identify metabolite quantitative trait loci for 44 urinary metabolites measured using proton NMR. The study included 996 children recruited as a part of the Human Early Life Exposome (HELIX) project who were genotyped using the Infinium Global Screening Array (GSA) MD version 1 (Illumina). Finally, Schlosser et al. [14] evaluated genetic associations of 1172 metabolites in 1627 participants from the German Chronic Kidney Disease (GCKD) study and the Study of Health in Pomerania Trend (SHIP-Trend). Metabolites were measured using mass spectrometry, and genotyping was conducted using Illumina Omni2.5Exome BeadChip arrays in the GCKD cohort, and Illumina HumanOmni2.5-Quad in the SHIP-Trend cohort.

### GWAS data processing

The extracted SNP-analyte association data included 1297 analyte associations to 1297 unique SNPs from the study by Nicholson et al.; 1 799 971 analyte associations to 608 024 unique SNPs from the study by Raffler et al.; 3 113 996 analyte associations with 780 302 unique SNPs from the study by Sinnott-Armstrong et al.; and 12 482 976 analyte associations with 283 704 unique SNPs from the study by Calvo-Serra et al.. Finally, due to the large number of associations in the study by Schlosser et al., only SNP-analyte associations matching the other studies were extracted, which amounted to 1 213 222 unique SNPs and 7 034 023 SNP-analyte associations. Following the removal of associations evaluated only by single studies, 14 179 833 SNP-analyte associations remained in the dataset, corresponding to 1 213 704 unique SNPs and 48 urinary metabolites.

### Meta-analysis of urinary metabolite GWAS studies

The sample size-based meta-analysis revealed 1226 SNPs with significant (P < 7.1 × 10^-9^) associations, of which 48 lead SNPs were identified following clumping for 14 urinary analytes. The respective analytes included alanine, 3-aminoisobutyrate, betaine, creatine, creatinine, formate, glycine, glycolate, histidine, 2-hydroxybutyrate, lysine, threonine, trimethylamine, and tyrosine. Details of the significant associations following clumping are shown in Supplementary Table S2. The findings included three novel significant associations that had not been reported previously, namely the association of rs4594899 to tyrosine (P = 6.6 × 10^-9^, N = 5447), rs1755609 to glycine (P = 3.3 × 10^-10^, N = 5447), and rs79053399 to 3-aminoisobutyrate (P = 6.9 × 10^-10^, N = 4656). The most significant SNP-analyte associations identified were those of rs37369 to 3-aminoisobutyrate (P ≤ 10^-308^, N = 6145) and rs2147896 to trimethylamine (P = 5.0 × 10^-165^, N = 1207), followed by the associations of rs163908 to 3-aminoisobutyrate (P = 2.6 × 10^-112^, N = 4975) and rs1154510 to 2-hydroxybutyrate (P = 1.8 × 10^-102^, N = 4806).

### Post-hoc analyses

The newly identified loci did not show any pleiotropic effects, as none of them were significantly associated with other phenotypic traits in the PhenoScanner database. Furthermore, a relatively large association cluster was identified on chromosome 17, as shown in Figure 2. The cluster contained significant SNP-metabolite associations to alanine, betaine, histidine, threonine, and tyrosine. A subsequent pathway analysis revealed aminoacyl-tRNA biosynthesis as a significant pathway involving four of these metabolites (P_chr17_ = 3.9 × 10^-6^).

**Fig 2.**
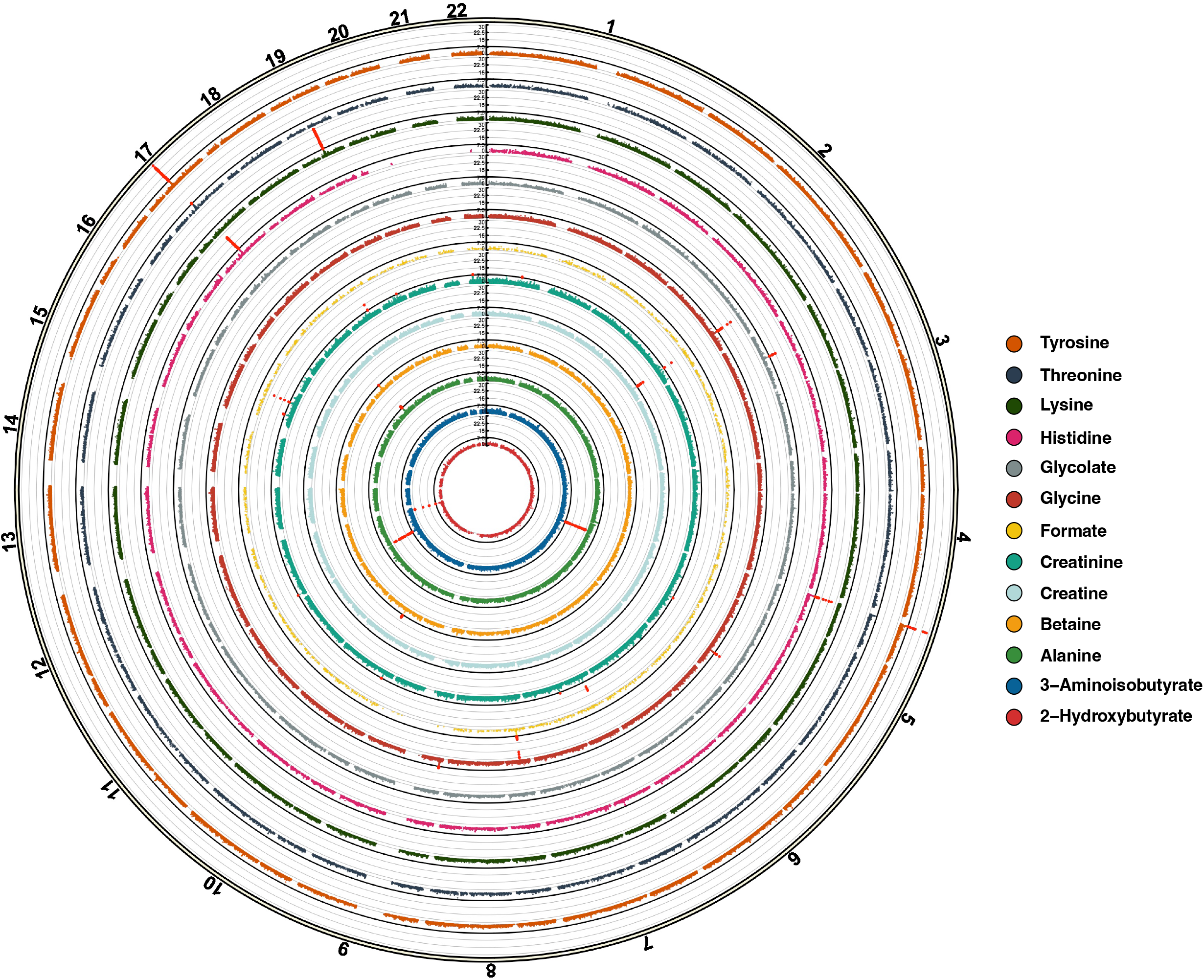
Circular Manhattan plot of all associations identified in the urinary metabolite GWAS meta-analysis. Trimethylamine was excluded from the plot due to a low number of associations. Chromosomes are indicated above the circle, numbered through 1-22. Red points on the plot, crossing the blue line, indicate significant associations (P < 7.1 x 10^-9^). Associations exceeding the threshold of P < 5 x 10^-30^ have been capped at the threshold for clarity of the plot.

## Discussion

The present study aimed to identify novel urinary metabolite loci through a comprehensive meta-analysis of urinary metabolite GWAS studies. To our knowledge, this meta-analysis represents the first such study of urinary metabolite GWASs conducted to date. Three different resources, including the EMBL-EBI GWAS Catalog, PubMed, and metabolomix.com, were searched to identify a set of relevant urinary metabolite GWAS studies, followed by a collection of extensive GWAS summary statistics data. A subsequent sample size-based meta-analysis revealed 48 significant SNP-analyte associations. The results confirmed previously reported associations and uncovered three novel urinary metabolite loci, namely rs4594899 for tyrosine, rs1755609 for glycine, and rs79053399 for 3-aminoisobutyrate. Additionally, a region on chromosome 17 was identified to be associated with the levels of multiple urine analytes. Upon further examination, a shared pathway for aminoacyl-tRNA biosynthesis was identified for most of the clustered associations.

Most of the newly discovered SNP-analyte associations are consistent with existing literature. The glycine locus rs1755609 is an intronic variant in the glycine decarboxylase (GLDC) gene region which encodes glycine dehydrogenase, also known as the P protein. This protein constitutes a central enzyme in the glycine cleavage system (GCS), where its primary role is to catalyze the decarboxylation of glycine [15]. Glycine is a major amino acid, primarily synthesized in the kidneys and the liver, and has significant roles in metabolic regulation, neurological function, and response to oxidative stress [16]. An excess of glycine in the body can lead to glycine encephalopathy, also known as nonketotic hyperglycinemia, a condition primarily caused by dysfunction in the GCS [16]. Additionally, imbalances of glycine and glycine-related compounds have been further linked to various neurobiological disorders such as bipolar disorder [17–19]. Our findings indicate that the genetic influences on the metabolic glycine degradation system extend to detectable changes in urinary glycine levels. This result could pave the way for future studies exploring the role of peripheral glycine in disease development and brain health.

The second of the identified metabolites with a novel genetic locus, 3-aminoisobutyrate, also known as beta-aminoisobutyric acid or BAIBA, is a product of thymine and valine metabolism. Considered an important player in carbohydrate and lipid metabolism, it is strongly upregulated by exercise, and therefore also associated with a lower risk of cardiometabolic diseases [20–23]. Its newly identified locus, rs79053399, is an intronic variant of the retinoic acid induced 14 (RAI14) gene which encodes the Rai14 protein, also known as ankycorbin. Rai14 is known to regulate mechanical properties of cells by contributing to membrane protein binding and shaping [24], and plays a role in actin organization and function. As such, it is involved in processes like cell motility, adhesion, polarity, and cytokinesis [25], and also acts as a key linking scaffold during micropinocytosis and cell migration [24]. While the mechanistic link between BAIBA and Rai14 is not clear, SNPs within the RAI14 gene have been previously linked to other metabolite alterations, such as those in creatinine [26] and uric acid levels [27].

This suggests that Rai14 could potentially regulate peripheral levels of multiple analytes in a non-specific manner. However, further research is required to elucidate this possible link.

The final of the newly identified loci, rs4594899, is a variant of the LINC02142 gene which encodes a long intergenic non-coding ribonucleic acid, or lincRNA. lincRNAs have numerous functions, including regulation of chromatin topology and neighboring genes, scaffolding and decoying for proteins and RNAs, as well as potentially contributing to micro-peptide production [28]. However, because the specific functions of LINC02142 remain unknown, its direct association to tyrosine is unclear. Past GWAS studies have found associations of SNPs within the LINC02142 gene to broad cognitive capabilities such as mathematical ability, educational attainment, and longitudinal attention [29, 30], addictive substance use involving caffeine and nicotine products [31, 32], and behavioral traits such as sedentary behavior and subjective wellbeing [33, 34]. Consistently, tyrosine is a metabolic precursor of the neurotransmitter dopamine [35], which is central to the reward mechanisms involved in the above-mentioned traits. Considering the close proximity of the LINC02142 gene with neighboring SNPs on chromosome 5, which are also associated with urinary tyrosine levels, and its regulatory function, it could be hypothesized that the product of the LINC02142 gene may act as a potential regulator of these loci. This, in turn, could affect dopamine synthesis and precipitate other LINC02142-associated traits. However, further research is needed to evaluate this hypothesis.

Among the significant associations confirmed by the present study, a notable cluster of five closely situated loci was observed on chromosome 17. These loci were linked with alanine, betaine, histidine, threonine, and tyrosine urine levels. Pathway analysis indicated that, except for betaine, these analytes are involved in the aminoacyl-tRNA biosynthesis pathway. Aminoacyl-tRNA synthesis is a process where amino acids are attached to the respective transfer-RNA molecules by aminoacyl-tRNA synthetases prior to translation [36, 37]. We hypothesise that the identified associations may converge on a common intermediary factor in the aminoacyl-tRNA synthesis pathway, possibly through modulating enzyme activity, thereby affecting the levels of multiple analytes. Alternatively, the findings could reflect underlying correlations among the urinary concentrations of tyrosine, threonine, and histidine, as all three analytes exhibit significant SNP associations in close proximity also elsewhere in the genome (chr 5). The present results highlight the need for further research to unravel the mechanisms behind the clustering of these associations.

Finally, our analysis provides valuable insights for enhancing future urinary metabolite GWAS studies. A pivotal observation from this research is the narrow focus of existing urinary metabolite GWAS studies, which predominantly target metabolites already known to be associated with specific diseases. This is exemplified by the fact that only 48 analytes were eligible for the current meta-analysis, as compared to nearly 4,000 compounds discovered in urine to date [38]. While such a focused approach has inherent value in disease-specific contexts, it severely constrains the general utility of these studies for other applications. In contrast, blood-based GWAS studies often rely on hypothesis-free metabolomic profiling which has proven to be broadly applicable, shedding light on a wide array of physiological and pathological states [2, 39, 40]. In contrast to blood samples, the non-invasive nature of collecting urinary metabolite readouts makes them promising candidates for broader research applications. Therefore, future studies should adopt a more generalist viewpoint, targeting a wide range of metabolites that can serve a multitude of research objectives. Furthermore, the accessibility of complete GWAS summary statistic data from the assessed studies was very limited, and many were not available even after contacting the authors. Therefore, open data access practices are critical for facilitating more rigorous meta-analyses and cross-disciplinary research.

The interpretation of our findings is subject to several limitations. First, in the absence of better alternatives, this meta-analysis utilized a sample size-based method to calculate statistical significance. This approach, while practical, is not ideal as it does not allow for the assessment of combined effect sizes and may lead to results being disproportionately affected by the largest of the included studies. Second, the number of meta-analyzed studies was low, and their ethnic composition was predominantly Caucasian, which may impact the robustness and generalizability of the findings. Third, the variations in participant demographics, such as young age or impaired kidney function, could have influenced urinary metabolite expression, potentially inflating or masking certain analyte-SNP associations. Finally, urinary analyte concentrations are inherently variable, both between and within individuals, due to factors such as diet, hydration, or activity levels [41, 42]. This could significantly influence the outcomes of the original GWAS studies included in our analysis, affecting the reliability of the findings.

## Conclusion

In conclusion, the present meta-analysis of five urinary metabolite GWAS studies identified three novel SNP-metabolite associations, including rs4594899 with tyrosine, rs79053399 with 3-aminoisobutyrate, and rs1755609 with glycine. The associations involving tyrosine and glycine are consistent with existing literature, while the potential link between 3-aminoisobutyrate and the protein Rai14 warrants further exploration. Our study highlights the necessity for an increased scope and larger sample size in future urinary metabolite GWAS studies, as well as the importance of open data access practices.

## Supporting information

**S1. Description of studies included in the meta-analysis**.

**S2. Results of the sample-size weighted meta-analysis of urinary metabolite GWAS studies**.

**Supplementary Data. Complete summary statistics of the meta-analysis**.

## Funding

This work was supported by the Stanley Medical Research Institute (grant number: O7R-1888) by grants to Sabine Bahn, and by the Oskar Huttunen Foundation grant to Jihan K. Zaki.

## Conflicts of Interest

Dr Tomasik has a patent pending for dried blood spot biomarkers for bipolar disorder.

Dr Bahn reported grants from Stanley Medical Research Institute and Psyomics during the conduct of the study; is a founder and shareholder in Psyomics; is Director of Psynova Neurotech outside the submitted work; and has a patent pending for dried blood spot biomarkers for bipolar disorder.

The other authors have no conflicts to declare.

## Author Contribution Statement

Conceptualization: SB, OAS, JT, JKZ; Methodology: SB, OAS, JT, JKZ; Data curation: JKZ; Data analysis: JKZ; Resources: SB, OAS; Writing - Original Draft: JKZ; Writing - Review & Editing: All co-authors; Supervision: SB, OAS, JT; Funding acquisition: SB, OAS, JKZ

